# Phytochemical Investigation of the Crude and Fractionated Extracts of two Nigerian Herbs, *Mitragyna Inermis* (Wild) and *Lawsonia Inermis* (Linn)

**DOI:** 10.1101/2021.06.02.446752

**Authors:** O.A. Onuh, M. Odugbo, O.O. Oladipo, I.W. Olobayotan

**Author notes:** **Corresponding Author: Olukemi, O.A., **.

## Abstract

Investigation was done to determine the phytochemicals present in the crude extracts and fractions of *Mitragyna inermis* (Wild) of the family, Rubiceae and *Lawsonia inermis* (Linn) of the family, Lythraceae- Loose - Strife using standard protocols to ascertain their bioactive molecules of pharmaceutical potentials. Successive extraction of the plant parts was carried out using hexane, ethyl acetate, methanol and water (aqueous). Eleven fractions were eluted from both plants using column and thin-layer chromatography. The study revealed the presence of alkaloids, flavonoids, saponins, phenols, tannins, steroids and carbohydrates in both plants while anthraquinone was present in only *L. inermis*. Quantitative study of some of the phytochemicals revealed various percentages ranging from 0.56% (flavonoids) to 21.58% (saponins). Alkaloids and phenols were present in more fractions (M6 – M11) of *M. inermis* extracts while tannins and anthraquinones were present in more fractions (L6 – L11) of *L. inermis* extracts than in other fractions. The phytochemicals present in these two plants could be responsible for their various therapeutic uses that have been reported. It is recommended that further research is done to isolate these bioactive molecules in their pure forms and evaluate the dosage required and toxicity levels.

## 1.0 Background of Study

Being naturally gifted by a suitable diverse range of climate from tropical to Savannah and fertile soil, Nigeria in West Africa possesses a rich flora of indigenous plants which are used in herbal medicine preparations to cure diseases and heal injuries (Ali, 2008; Sofowora, 2008; Soladoye *et al*., 2010). *Mitragyna inermis* (Willd) and *Lawsonia inermis* (Linn) are two of such medicinal plants with great potentials. Their ethnomedical importance has been reported in various indigenous systems of folk medicine and scientific documents (Quedraogo *et al*., 2004; Ali, 2008; Sofowora, 2008). These plants have been used widely in the treatments of diseases such as syphilis, typhoid fever, jaundice, Acquired Immune Deficiency Syndrome (AIDS), skin infections (Chaudhary *et al*., 2010). The powder/paste forms of the root, stem and leaf extracts of the two plants have also been reported to be used in treatment of asthma, pains, blood, skin and lung diseases, cataract and malaria (Mishana *et al*., 2000; Simplice *et al*., 2011).The following phytochemicals have been associated with the pharmacological activities of the selected plants: saponin, tannin, phenol, terpenoids, carbohydrate, alkaloids, glycosides, flavonoids, sterols (Elmanama *et al*., 2011; Muhammad and Muhammad, 2005).

Plants and their bioactive constituents have proved valuable in the treatment of a wide spectrum of health challenges, yet prospects for discovery of new chemotherapeutic agents from plant sources still abound. People have attested to the healing potentials of herbal plants in diverse pathological conditions (Ode *et al*., 2011). Despite appreciable progress made in the management of various diseases using the conventional therapeutic schemes, the search for plant based products for the control of diseases still continues. Since indigenous herbal medication has been documented as one of the main source of immediate help for populations that lack access to formal health resources in remote and rural environs, it is important to understand indigenous plants, especially their antimicrobial properties, etiology and treatment in order to improve health status in developing nations. Study after study has shown that the effect produced by extracts of whole plants is less than the effect produced when isolated purified constituents of the plant is administered. Hence, the extraction of bioactive compounds from plants is the first step in the utilization of its phytochemicals in herbal preparations. Herbal formulations are getting more important in the treatment of diseases due to the adverse effects of the synthetic drugs (Prassad *et al*. 2012). Reports are available individually on the antimicrobial activity of crude extracts of the selected plants, but there is little or no information of the safety (toxicology) and fractionation of the crude extracts into phyto compounds. Thus in order to provide a scientific justification for the utilization of these plants as therapeutic agents, this study was done to ascertain the phytochemicals present in their extracts.

## 2.0 Methods

### 2.1 Extracts Preparation

The crude plant extracts were prepared by the methods of Aboh *et al.* (2014). The samples were extracted using soxhlet apparatus on a rotary shaker with n-hexane, ethyl acetate, methanol (70%) and sterile distilled water successively. Five hundred grammes of each pulverized parts was extracted using n-hexane for 48h.The residue was dried in oven, while the collected extract was concentrated through evaporation under reduced pressure, packed and kept at 4 ◻C for further biological investigations. The dried residue was further extracted with ethyl acetate for another 48h. The extract was also concentrated as above while the residue was dried and extracted with methanol, concentrated as above and finally extracted with sterile distilled water. All the crude extracts were filtered and concentrated through evaporation under reduced pressure then transferred into sterile sample bottles, labeled appropriately and kept in refrigerator for further use.

### 2.2 Sterilization of the plant extracts

The extracted parts above were filtered using the membrane filtration system as described by (Sultana, 2007). The membranes are held in holders, supported on a frame. Fluids are made to transverse membranes by negative pressure. The filter membrane disc used is made of cellulose ester having a nominal average pore diameter of 30 micron (0.30mm). The membrane was held firmly in a filtration unit which consists of a supporting base for the membrane, a receptacle for the fluids to be filtered, and a collecting reservoir for the filtered fluid. This was carried out under aseptic condition.

### 2.3 Fractionation of the crude methanolic extracts of the plants

Fractionation of the crude methanolic leaf extracts of *L. inermis* and *M. inermis* were carried out in two parts: Column and Thin Layer Chromatography. The protocols of Nwodo *et al*. (2010), Ode *et al*. (2011) and Adefuye and Ndip (2013) with slight modifications were employed.

#### 2.3.1. Column chromatography

The crude leaf extracts of *L. inermis* and *M. inermis* were subjected to column chromatography to separate them into component fractions. The stationary phase (absorbent) used is column graded silica gel 60G (MERCK) while combinations of hexane, ethyl acetate and methanol were used as mobile phase (solvent system). In the setting up of the column chromatography, glass column of internal diameter 80mm and length 100cm (Quickfit, England) was used. The lower part of the glass column was plugged with glass wool with the aid of glass rod. Sand bed was placed over the glass wool. The sand bed served to give a flat base to the column of the absorbent. The wet packing method was used in preparing the silica gel column. 25g of silica gel (200-425 mesh particle, size Å pore) was wet packed with 250ml hexane solvent system. The slurry of the silica gel and hexane was poured down into the column carefully. The tap of the column was left open to allow free flow of solvent into a conical flask below. The set up was seen to be in order when the solvent drained freely without carrying the silica gel, sand, glass or wool into the tap. At the end of the packing process, the tap was locked. The column was allowed to stabilize for about 24h. A slurry of the crude extract was prepared in a ceramic mortar by adsorbing 5g of the extract to 10g of silica gel in 10ml of methanol. The slurry was gently loaded onto the packed column. The column was then eluted with solvent systems (mobile phase) gradually in order of increasing polarity using hexane, ethyl acetate and methanol, at a ratio of 2:1 v/v.

The following ratios of the solvent combinations were sequentially used in the elution process:

Hexane: ethyl acetate 100:0; 80:20; 60:40; 40:60; 20:80; ethyl acetate: methanol: 100:00; 80:20; 60:40; 40:60; 20:80; 0:100.

The above solvent systems (mobile phase) were continuously poured from the edge of the column with the aid of a dropper. The bottom outlet of the column was opened allowing eluent to flow through the column. As the eluent passed down the column, the compound fraction moved down the column. The separated fractions flowed out of the column where the different eluents were collected in properly labeled bottles. TLC analysis was carried out on fractions before they were evaporated over water bath. The weights of the dry fraction were recorded.

#### 2.3.2 Thin Layer Chromatography

Analytical TLC plates were prepared by pouring homogenous silica gel (60, F_254_, MERCK) slurry into aluminum plates by spreading technique. The silica gel layer was adjusted to 0.25mm thickness. The coated plates were allowed to dry in air and activated by heating in hot air oven at 100-150◻C for 1h.

#### 2.3.3 Preparation of the development tank (mobile phase)

Solvent system used was hexane and ethyl acetate at the ratio of 4:1. v/v, then ethyl acetate (less polar) and methanol (more polar). The fractions obtained from the column chromatography were spotted with the help of capillary tube washed in acetone. Each fraction was applied as a single spot in a row along one side of the chromo plate, 2cm from the edge and 1.5cm from edge (known as the origin). The spotted chromo plate was placed at an angle of 45◻ in the development tank containing the solvent system, covering the bottom of the plate by the solvent up to nearly 1cm.The solvent front was marked on the plate immediately after removing it from the chamber and allowed to dry. The mobile phase was not allowed to reach the end of the stationary phase.

The plate was visualized and dried by hot air dryer. The plate was then viewed under daylight and UV light at 302nm and 365nm respectively. The plate was further exposed to iodine fumes in a chamber and finally sprayed with freshly prepared Vanillin reagent (0.16g Vanillin powder + 14ml of methanol + 0.5ml concentrated sulphuric acid. The plate was carefully heated at 105◻C for optimal colour development. Characterization of the different compounds identified was done by calculating Rf values. (Adefuye and Ndip, 2013). The Rf value is:

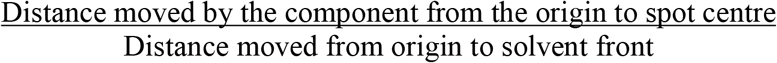

Where solvent is stationary.

The fractions showing similar TLC mobility and band (that is, same RF) were pooled together. The fractions were kept at 4◻C in the refridgerator for further bioassay tests to confirm their biological activity.

#### 2.3.4 Determination of minimum inhibitory concentration (MIC) of eluted fractions

The MIC was determined by using the micro-dilution method as described by Adefuye and Ndip (2013) and Aboh *et al.,* (2014) with slight modifications. This assay was performed using round bottom 96-well microtitre plate. Two-fold serial dilutions of the eluted fractions were prepared in the test wells starting with 50mg/ml stock. The dilutions were 25mg/ml, 12.5mg/ml, 6.25 mg/ml, 3.125mg/ml, 1.565mg/ml. The bacterial strains were purified by standard bacteriological methods (Cheesebrough, 1984, CCLS, 2002). The bacterial strains were standardized to 0.5 MacFarland standard using Nephelometer to give approximately 1.0 × 10^6^ cfu/ml. The standardized cultures were maintained in sterile Muller Hinton agar. Eighteenth-hour broth culture of the *Candida albicans* was suspended into sterile Sabouraud dextrose liquid medium. It was standardized according to Clinical Laboratory Standards Institutes (CLSI, 2002) by gradually adding normal saline to compare its turbidity to McFarland standard of 0.5 which is approximately 1.0 × 10^6^ cfu/ml using Nephlometer.

Control wells were prepared using Ciprofloxacin 20mg/ml as positive for bacteria. Dimethyl sulfoxide (DMSO) was used as negative control. The plates were then sealed with parafilm and incubated at 37^°^C for 24hrs for bacteria and yeast. After incubation, 40μL of 0.2mg/ml of Tetrazonium dye (p-iodonitrotetrazoluim violet) was added to well and incubated for an hour, after which the uninhibited organisms would have converted the dye from blue to pink in the well. The MIC was defined as the minimum concentration at which growth was inhibited with no visible change. Sterility test was performed to verify whether the broth used in the assay was contaminated before test procedures. 50μL of broth was dispensed into a well, without both extract and inoculums.

### 2.4 Preliminary Phytochemical Screening

The phytochemical screening for the crude extracts was carried out using standard protocols as described by Sofowora (2008), Trease and Evans (1999) as modified by Aboh *et al*., (2014). The following phytochemicals were screened for alkaloids, cardiac glycosides, tannins, flavonoids, phenols, saponins, anthraquinones, sterols and terpenes.

#### 2.4.1 Test for tannins

About 10 mg of the plant extract was taken and was dissolved in 45 % of ethanol. The sterile test tubes were boiled for five minutes and 1 mL of 15% ferric chloride solution was added to each. The appearance of greenish to black colour confirms the presence of tannins in the plant extracts.

#### 2.4.2 Test for flavonoids

##### Magnesium chips or Shinoda test

The powdered plant (0.5 gm) was extracted in ethanol by boiling on a water bath for 5 min, filtered and cooled. A small quantity of Magnesium chips was added to the filtrate and few drops of concentrated HCl were added down the side of the test-tube. The colour change was noted.

##### Zinc chips test

A small quantity of zinc chips was added to the powdered plant and few drops of concentrated HCl were added down the side of the test tube. The colour change was noted.

#### 2.4.3 Test for resins

Fifteen milliliters of plant were prepared using 0.1 g of the powdered plant and filtered into a test-tube. An equal volume of copper acetate solution was added and shaken vigorously then allowed to separate. The colour change was noted.

The powdered plant (0.5g) was dissolved in acetic anhydride and one drop of concentrated sulphuric acid was added. The colour change was noted.

#### 2.4.4 Test for Steroids

##### Liebermann–Burchard test

Powdered plant suspension (1mL) was treated with chloroform, acetic anhydride and drops of H_2_SO_4_ was added and observed appearance of dark reddish green colour confirmed the presence of steroids in the plant extract.

A fraction of plant was treated with ethanol and H_2_SO_4_ and observed for the formation of violet blue or green colour.

#### 2.4.5 Test for Phenols

##### Ferric chloride test

A fraction of plant was treated with 5 % ferric chloride and observed for formation of deep blue or black colour.

##### Liebermann’s test

The plant was heated with sodium nitrite, added H_2_SO_4_ solution diluted with water and excess of dilute NaOH was added and observed for the formation of deep red or green or blue colour.

#### 2.4.6 Test for Anthraquinones

##### Borntrager’s test

About 50 mg of powdered extract was heated with 10% ferric chloride solution and 1ml concentrated HCl. The extract was cooled, filtered and the filtrate was shaken with diethyl ether. The ether extract was further extracted with strong ammonia; pink or deep red colourations of aqueous layer indicate the presence of anthraquinone.

#### 2.4.7 Test for Alkaloids

##### Wagner’s test

A fraction (10 mg) of the plant extract was treated with Wagner’s test reagent [1.27 g of iodine and 2 g of potassium iodide in 100 mL of water] and observed for the formation of reddish brown colour precipitate, which confirmed the presence of alkaloids in the plant extract.

#### 2.4.8 Test for Terpene

##### Salkowski test

Five mL of each plant mixture was mixed in 2 mL of chloroform, and concentrated H_2_SO_4_ (3 mL) was carefully added to form a layer. A reddish brown colouration of the inter face was formed to show positive results for the presence of terpenoids.

#### 2.4.9 Test for saponin

Ten mg of the powdered extract sample was boiled (80-100◻) in 20 mL of distilled water in a water bath and filtered. 10mL of the filtrate was mixed with 5 mL of distilled water and shaken vigorously for a stable persistent froth. Formation of foam (froth) on top of the test tube shows the presence of saponins. Three (3) drops of olive oil was added to the mixture and shaken vigorously, then observed for the formation of emulsion.

### 2.5 Quantitative Determination of Phytochemical Constituents of the Plant Samples

The standard protocols described by Soladoye and Chukwuma (2012) were employed.

#### 2.5.1 Determination of Alkaloids

This was done by the alkaline precipitation gravimetric method described by Harborne (1998). A measured weight of the sample was dispersed in 10% acetic acid solution in ethanol to form a ratio of 1:10 (10%). The mixture was allowed to stand for 4h at 28◻C. It was later filtered via Whatman No. 42 grade of filter paper. The filtrate was concentrated to one quarter of its original volume by evaporation and treated with drop wise addition of conc. aqueous NH_4_OH until the alkaloid was precipitated. The alkaloid precipitated was received in a weighed filter paper, washed with 1% ammonia solution dried in the oven at 80◻C. Alkaloid content was calculated and expressed as a percentage of the weight of sample analyzed.

#### 2.5.2 Determination of Flavonoids

This was also determined according to the method outlined by Harbone, (1998). 5gram of the sample was boiled in 50ml of 2M HCl solution for 30min under reflux. It was allowed to cool and then filtered through Whatman No 42 filter paper. A measured volume of the extract was treated with equal volume of ethyl acetate starting with a drop. The flavonoid precipitated was recovered by filtration using weighed filter paper. The resulting weight difference gave the weight of flavonoid in the sample.

#### 2.5.3 Determination of Tannins

Swain’s method was used for the determination of tannin contents of the powdered sample. 0.2 g of finely ground sample was measured into a 50 mL beaker. 20 mL of 50% methanol was added and covered with paraffin and placed in a water bath at 77-80◻C for 1 h and stirred with a glass rod to prevent lumping. The extract was quantitatively filtered using a double layered Whatman No.1 filter paper into a 100 mL volumetric flask using 50% methanol to rinse. This was made up to mark with distilled water and thoroughly mixed. 1 mL of sample extract was pipetted into 50 mL volumetric flask, 20 mL distilled water, 2.5 mL Folin-Denis reagent and 10 mL of 17% Na_2_CO_3_ were added and mixed properly. The mixture was made up to mark with distilled water, mixed well and allowed to stand for 20 min when a bluish-green colouration developed. Standard Tannic Acid solutions of range 0-10 ppm were treated similarly as 1 mL of sample above. The absorbance of the Tannic Acid Standard solutions as well as samples were read after colour development on a Spectronic 21D Spectrophotometer at a wavelength of 760nm. Percentage of tannin was calculated.

#### 2.5.4 Determination of Saponin

The Spectrophotometric method described by Brunner was used for saponin analysis. 1 g of finely ground sample was weighed into a 250 mL beaker and 100 ml Isobutyl alcohol was added. The mixture was shaken on a UDY shaker for 5 h to ensure uniform mixing. Thereafter, the mixture was filtered through a Whatman No. 1 filter paper into a 100 mL beaker and 20 mL of40% saturated solution of Magnesium carbonate added. The mixture obtained with saturated MgCO_3_ was again filtered through a Whatman No. 1 filter paper to obtain a clear colourless solution. 1 mL of the colourless solution was pipetted into 50 mL volumetric flask and 2 mL of 5%FeCl_3_ solution was added and made up to mark with distilled water. It was allowed to stand for30 min for blood red colour to develop. 0-10 ppm standard saponin solutions were prepared from saponin stock solution. The standard solutions were treated similarly with 2 mL of 5% FeCl_3_solution as done for 1 mL sample 3 above. The absorbencies of the sample as well as standard saponin solutions were seen on Spectrophotometer at a wavelength of 380 nm. The percentage saponin was also calculated.

#### 2.5.5 Determination of Anthraquinone contents

About 50 mg of the fine powder sample was soaked in 50 mL of distilled water for 16 hours. This suspension was heated in water bath at 70◻C for one hour. After the suspension was cooled, 50 mL of 50% methanol was added to it and then filtered. The clear solution was measured by spectrophotometer at a wavelength of 450nm and compared with a standard solution containing1mg/100mL alizarin and 1mg/100mL purpurin with the absorption-maximum 450nm.

#### 2.5.6 Determination of Steroids

The steroid content of the leaves of the plants was determined using the method described by Harborne, (1998).5g of the powdered sample was hydrolysed by boiling in 50 mL hydrochloric acid solution for about 30minutes. It was filtered using Whatman filter paper (No. 42) the filtrate was transferred to a separating funnel. Equal volume of ethyl acetate was added to it, mixed well and allowed separate into two layers. The ethyl acetate layer (extract) recovered, while the aqueous layer was discarded. The extract was dried at 100◻C for 5minutes in a steam bath. It was then heated with concentrated amyl alcohol to extract the steroid. The mixture becomes turbid and a reweighed Whatman filter paper (No. 42) was used to filter the mixture properly. The dry extract was then cooled in a dessicator and reweighed. The process was repeated two more times and an average was obtained.

The concentration of steroid was determined and expressed as a percentage thus % Steroid = W2 –W1 × 100 Wt of sample 1

Where,

W1= weight of filter paper.
W2 = weight of filter paper + steroid

## 3.0 Result

All the fractions were further subjected to thin layer chromatography. Table 2 shows the details of the Thin Layer Chromatography profiles of the leaf extracts of *Lawsonia inermis.* The weight and percentage yield of the fractions of both plants and the retardation factor (Rf) values are also shown in the above tables. Fractions of *L. inermis* with same Rf values were pooled together to give the total of 6 fractions. They are:

LT1- fraction LT1
LT2- combination of fractions 2 and 3
LT3- fraction 4
LT4- fraction 5
LT5- combination of fractions 6 and 7
LT6- combination of fractions 8, 9, 10, 11

**Table 1:** Details of Fractions from Column Chromatography of Crude Methanolic Extract of *Lawsonia inermis* (L) Leaf.

| Fractions | Eluent<br>(Solvent System) | Ratio<br>V/V | Colour of Fraction |
| --- | --- | --- | --- |
| LC1 | Hex | 100 | Grey |
| LC2 | Hex : EA | 80 : 20 | Greenish |
| LC3 | Hex : EA | 60 : 40 | Dark green |
| LC4 | Hex : EA | 40 : 60 | Dark green |
| LC5 | EA | 20 : 80 | Light brown |
| LC6 | Hex : EA | 0 :100 | Brownish |
| LC7 | EA : Met | 80 : 20 | Brownish |
| LC8 | EA : Met | 60 : 40 | Reddish brown |
| LC9 | EA : Met | 40 : 60 | Reddish brown |
| LC10 | EA : Met | 20 :80 | Reddish brown |
| LC11 | Met : | 100 | Reddish brown |

Key: Hex = Hexane
EA = Ethyl Acetate
Met = Methanol
LC = *Lawsonia inermis* fraction from Column chromatography

**Table 2:** Details of Thin Layer Chromatography of Methanolic Extract of *Lawsonia inermis* Leaf.

| Eluted Fractions | Weight of Fraction (g) | Percentage (%) Yield | Rf Value |
| --- | --- | --- | --- |
| LT- 1 | 0.065 | 1.63 | 0.19 |
| LT- 2 | 0.213 | 5.33 | 0.34 |
| LT- 3 | 0.212 | 5.30 | 0.45 |
| LT- 4 | 0.302 | 7.56 | 0.61 |
| LT-5 | 0.944 | 23.59 | 0.71 |
| LT- 6 | 1.625 | 40.64 | 0.85 |

**Key:** Mobile System/ Solvent (Eluent) HEM = Hexane Ethyl Methanol Ratio of 4: 4: 1 (v/v) LT = *Lawsonia inermis* fraction from TLC

All the fractions were further subjected to thin layer chromatography. Table 4 shows the details of the Thin Layer Chromatography profiles of the leaf extracts of *Mitragyna inermis.* The weight and percentage yield of the fractions of the plant and the retardation factor (Rf) values are also shown in the above tables. Fractions with same Rf values for *Mitragyna inermis* were pooled together to give a total of 6 fractions. They are:

MT1- fraction LT1
MT2- combination of fractions 2 and 3
MT3- fraction 4
MT4- fraction 5
MT5- combination of fractions 6 and 7
MT6- combination of fractions 8, 9, 10, 11.

**Table 3:** Details of Fractions from Column Chromatography of Crude Methanolic Extract of *Mitragyna inermis* Leaf.

| FRACTIONS | ELUENT<br>(SOLVENT SYSTEM) | RATIO<br>V/V | COLOUR OF<br>FRACTION |
| --- | --- | --- | --- |
| MC1 | Hex | 100 | No |
| MC2 | Hex : EA | 80 : 20 | Grayish |
| MC3 | Hex : EA | 60 : 40 | Greenish |
| MC4 | Hex : EA | 40 : 60 | Dark green |
| MC5 | EA | 20 : 80 | Yellowish brown |
| MC6 | Hex : EA | 0 :100 | Yellowish brown |
| MC7 | EA : Met | 80 : 20 | Brownish |
| MC8 | EA : Met | 60 : 40 | Reddish |
| MC9 | EA : Met | 40 : 60 | Reddish |
| MC10 | EA : Met | 20 :80 | Reddish |
| MC11 | Met : | 100 | Reddish brown |

**Key:** Hex = Hexane
EA = Ethyl Acetate
Met = Methanol
MC = *Mitragyna inermis* fraction from column chromatography

**Table 4:** Details of Thin Layer Chromatography of Methanolic Extract of *Mitragyna inermis* Leaf.

| Eluted Fractions | Weight of Fraction (g) | Percentage (%) Yield | Rf Value |
| --- | --- | --- | --- |
| MT- 1 | 0.184 | 3.70 | 0.42 |
| MT- 2 | 0.235 | 4.71 | 0.56 |
| MT- 3 | 0.246 | 4.92 | 0.66 |
| MT- 4 | 0.237 | 4.74 | 0.78 |
| MT- 5 | 1.091 | 21.83 | 0.7 |
| MT- 6 | 2.611 | 52.22 | 0.89 |

**Note:** Mobile System/ Solvent (Eluent) HEM = Hexane Ethyl Methanol Ratio of 4: 4: 1 (v/v) MT = *Mitragyna inermis* fraction from TLC

### 3.1 Phytochemical Profile of the Crude Extracts

#### 3.1.1 Phytochemical Screening of the Crude Extracts of *Lawsonia inermis*

The phytochemical screening of the root, stem and leaf crude extracts of *Lawsonia inermis* carried out revealed the presence of some phytochemicals (Table 5). The hexane extract of the root revealed the presence of alkaloids, cardiac glycosides, saponins and resins. While stem extract revealed in addition to the ones in the root, flavonoids sterols and phenols. The hexane leaf extracts revealed nine phytochemicals namely alkaloids, cardiac glycosides, anthraquinones, terpenes, tannins, saponins, flavonoid, phenols and resins. Sterols were not detected.

**Table 5:**
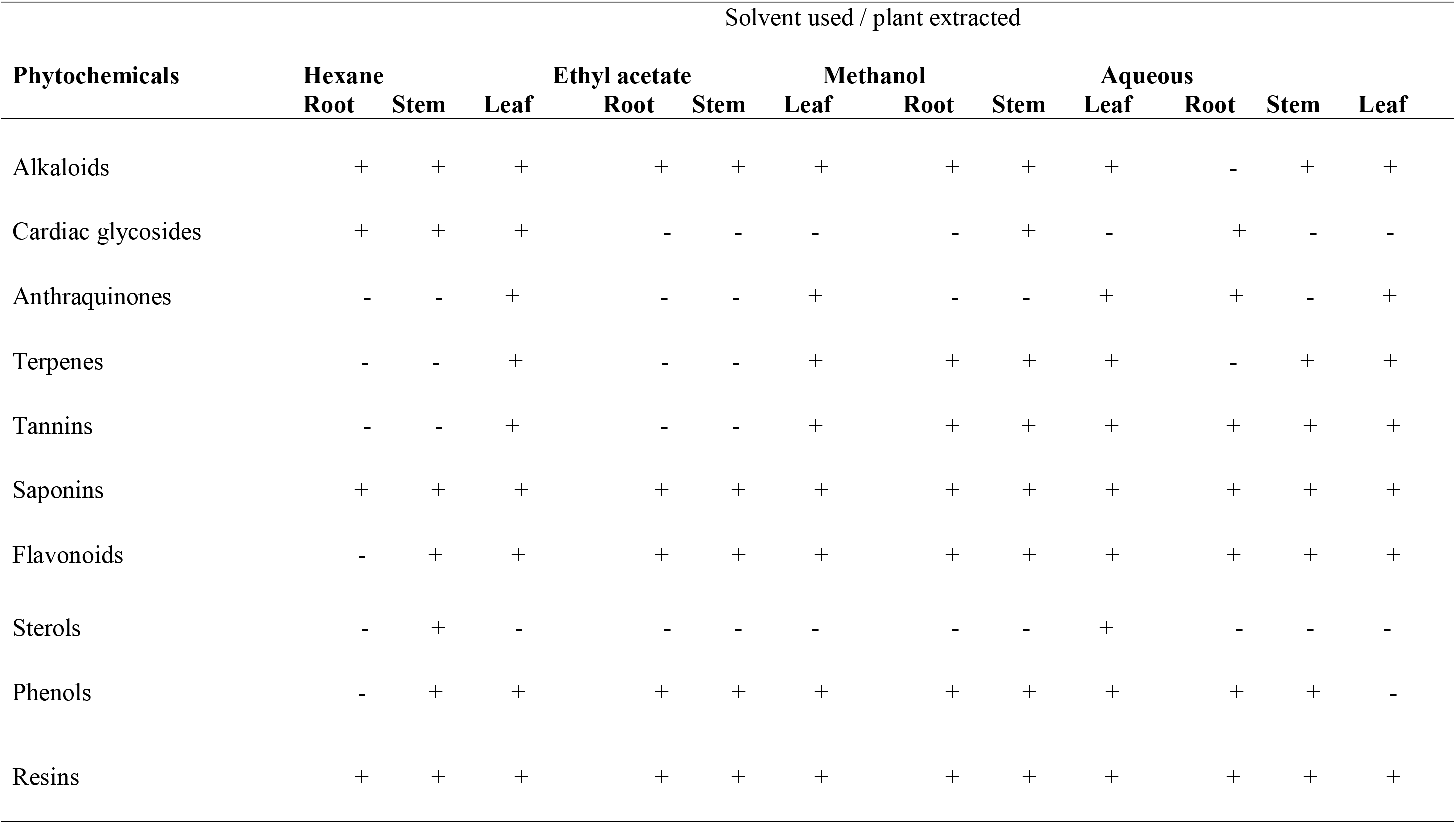
Phytochemical Screening of crude extracts of *Lawsonia inermis*.

The ethyl acetate root extract showed five (5) phytochemicals namely alkaloids, saponins, flavonoids, phenols, resins. The stem extract also showed a similar pattern of five phytochemicals present in the root. However, the leaf extract showed eight phytochemicals. Cardiac glycosides and sterols were absent in all the three aerial parts screened.

The methanol extract, showed a pattern of phytochemicals that was slightly different. All the phytochemicals screened with the exception of cardiac glycosides were present in the root, stem and leaf. The leaf showed the presence of alkaloids, anthraquinones, tannins, saponins, flavonoids, sterols, phenols and carbohydrates. Anthraquinones was not detected in the root and stem extract.

In the aqueous extract, sterols were absent in the root, stem and leaf. Seven phytochemicals were revealed in the root and the stem. The aqueous leaf extract showed the presence of alkaloids, anthraquinones, terpenes, tannins, saponins, flavonoids and resins. Tannins, saponins and flavonoids were present in all the aerial parts of *Lawsonia inermis*.

The percentage concentration (quantitative profile) of alkaloids, flavonoids, steroids, anthraquinones and tannins are shown in Table 6.

**Table 6:**
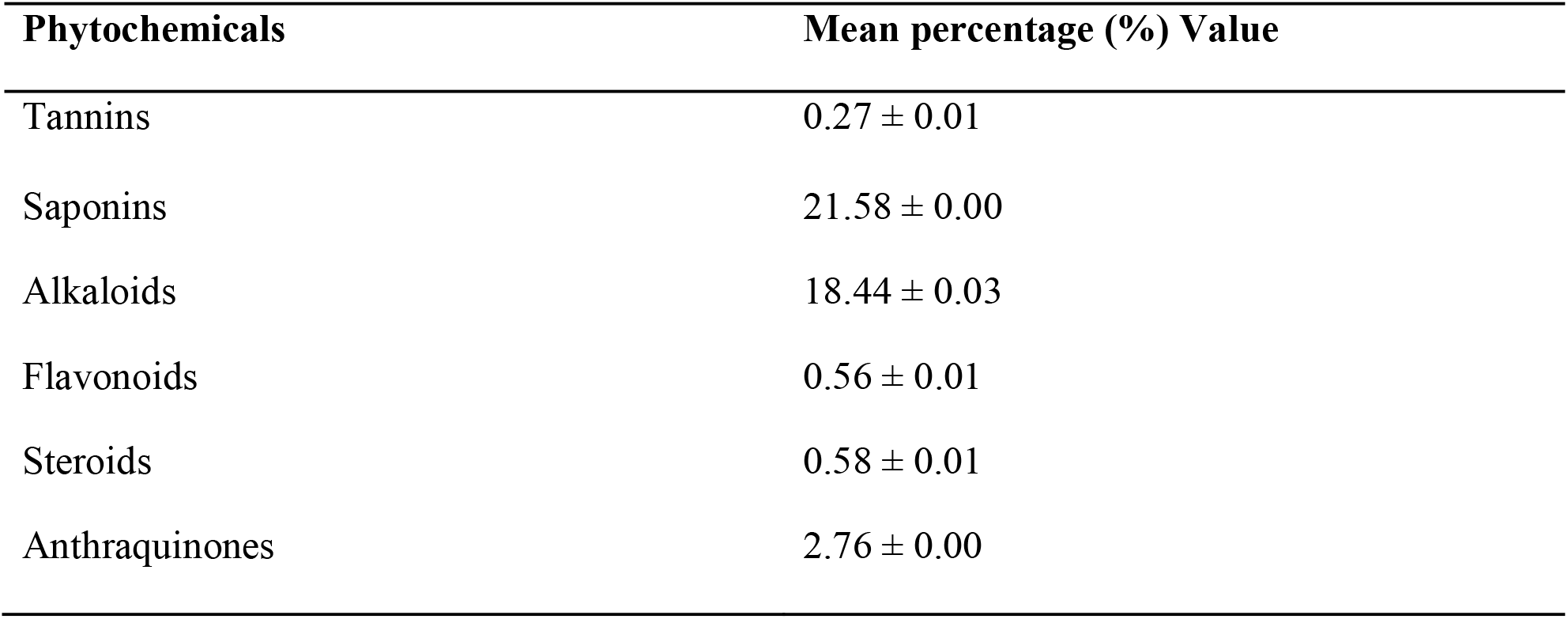
Quantitative Phytochemical Content of the methanol leaf extract of *L. inermis* (mean percentage - %)

| Phytochemicals | Mean percentage (%) Value |
| --- | --- |
| Tannins | $0.27 \pm 0.01$ |
| Saponins | $21.58 \pm 0.00$ |
| Alkaloids | $18.44 \pm 0.03$ |
| Flavonoids | $0.56 \pm 0.01$ |
| Steroids | $0.58 \pm 0.01$ |
| Anthraquinones | $2.76 \pm 0.00$ |

#### 3.1.2 Qualitative Phytochemical content of *Lawsonia inermis* fractions

Methanol extract of *Lawsonia inermis* was fractionated. Eleven (11) fractions (L1-L11) were eluted. The following phytochemicals were revealed in the eleven fractions: alkaloids, tannins, saponin, flavonoids, terpenes, cardiac glycosides, sterols, phenols, anthraquinones and resins. This is a confirmation of the phytochemicals that were present in the crude extracts. The details of the result are shown in Table 7.

**Table 7:**
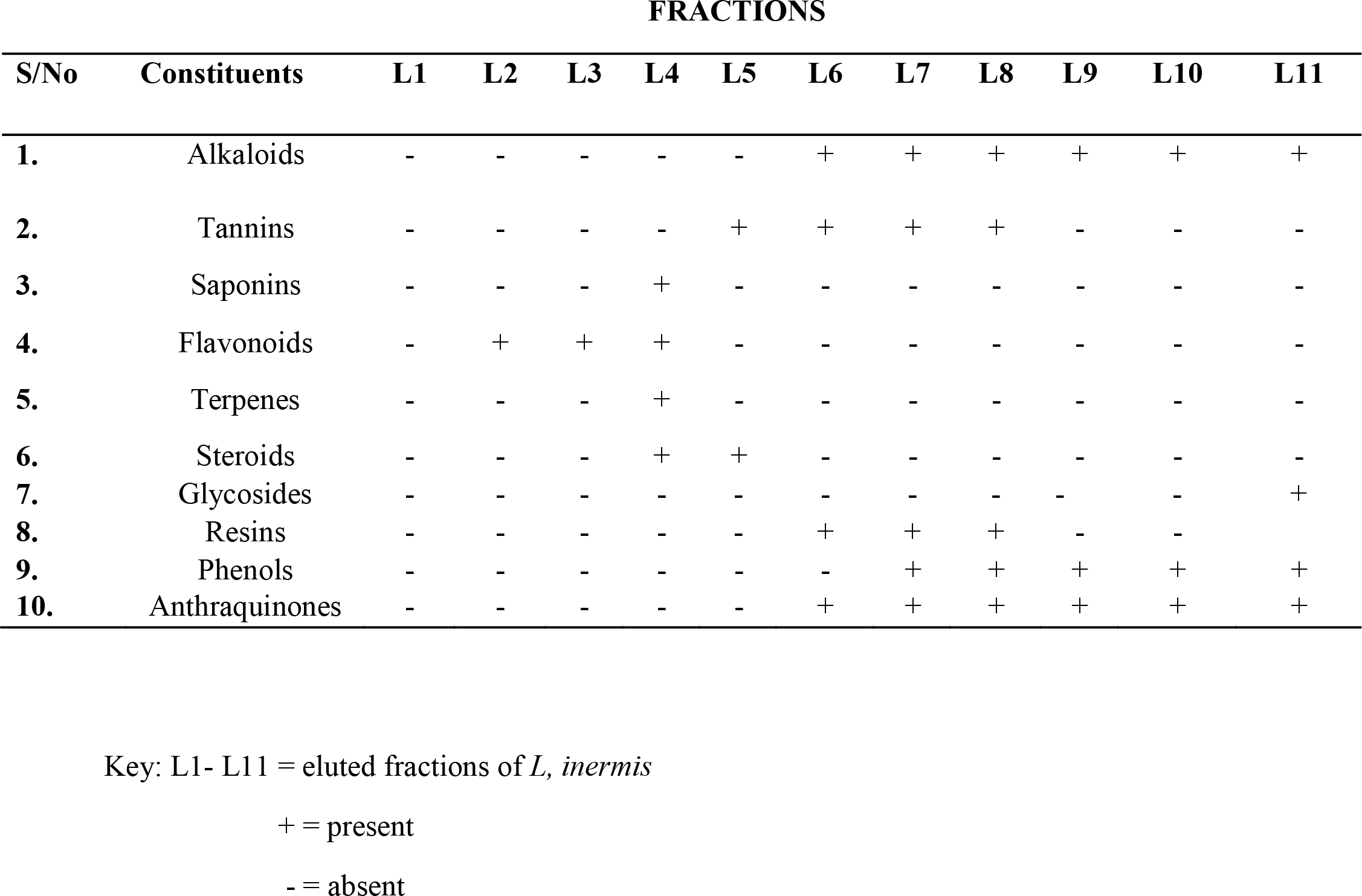
Qualitative Phytochemical Content of *Lawsonia inermis* fractions FRACTIONS.

Alkaloids were present in fractions L6, L7, L8, L9, L10, and L11. Tannins were present in only fraction L5-L8. Terpenes and Saponins were present in only fraction L4, while cardiac glycosides were present in only fraction L11.

#### 3.1.3 Phytochemical screening of the Extracts of *Mitragyna inermis*

The phytochemical screening of the root, stem and leaf crude extracts of *Mitragyna inermis* carried out revealed the presence of the following phytochemicals: alkaloids, terpenes, tannins, saponins, flavonoids, sterols, phenols and resins.

Anthraquinones and cardiac glycosides were absent in all the plant parts screened. Alkaloids terpenes, tannins and carbohydrates were present in all the solvents used on the plant part that is the root, stem, leaf. Saponin, flavonoid and phenol were present in the leaf extracts of hexane, ethyl acetate, methanol and aqueous. The details are shown in Table 8.

**Table 8:** Phytochemical Screening of crude extracts of *Mitragyna inermis*.

| Phytochemicals | Solvent used / plant extracted |  |  |  |  |  |  |  |  |  |  |  |
| --- | --- | --- | --- | --- | --- | --- | --- | --- | --- | --- | --- | --- |
|  | Hexane |  |  | Ethyl acetate |  |  | Methanol |  |  | Aqueous |  |  |
|  | Root | Stem | Leaf | Root | Stem | Leaf | Root | Stem | Leaf | Root | Stem | Leaf |
| Alkaloids | + | + | + | + | + | + | + | + | + | + | + | + |
| Cardiac glycosides | - | - | - | - | - | - | - | - | - | - | - | - |
| Anthraquinones | - | - | - | - | - | - | - | - | - | - | - | - |
| Terpenes | + | + | + | + | + | + | + | + | + | + | + | + |
| Tannins | + | + | + | + | + | + | + | + | + | + | + | + |
| Saponins | - | - | + | - | - | + | - | - | + | - | - | + |
| Flavonoids | - | - | + | - | - | + | - | - | + | - | - | + |
| Sterols | - | - | + | - | - | + | - | - | + | - | - | - |
| Phenols | - | - | + | - | + | + | + | + | + | + | + | + |
| Resins | + | + | + | + | + | + | + | + | + | + | + | + |

Key: + = Positive; - =Absent

#### 3.1.4 Qualitative Phytochemical Contents of *Mitragyna inermis* fractions

Phytochemical analysis of the fractionated methanol extract of *Mitragyna inermis* leaf revealed the presence of the following; alkaloids, tannins, flavonoids, terpenes, phenols, saponins, sterols, resins. Eleven fractions were eluted (M1-M11). No single fraction contained all the phytochemicals screened, but all the phytochemicals in the crude were present in the fractions. Table 9 shows the details of the result obtained. Anthraquinone was absent in all the fractions.

**Table 9:** Qualitative Phytochemical Contents of *Mitragyna inermis* Fractions.

| S/No | Constituents | M1 | M2 | M3 | M4 | M5 | M6 | M7 | M8 | M9 | M10 | M11 |
| --- | --- | --- | --- | --- | --- | --- | --- | --- | --- | --- | --- | --- |
| 1. | Alkaloids | - | - | - | - | + | + | + | - | - | - | - |
| 2. | Tannins | - | - | - | - | - | + | + | + | + | + | + |
| 3. | Saponins | - | - | - | - | - | - | - | - | - | - | - |
| 4. | Flavonoids | - | - | - | + | + | - | - | - | - | - | - |
| 5. | Terpenes | - | - | - | - | + | + | + | - | - | - | - |
| 6. | Steroids | - | - | - | - | - | - | - | - | - | - | - |
| 7. | Glycosides | - | - | - | - | - | - | - | - | - | - | - |
| 8. | Resins | - | - | - | - | - | - | - | - | - | - | - |
| 9. | Phenols | - | - | - | - | - | + | + | + | + | + | + |
| 10. | Anthraquinones | - | - | - | - | - | - | - | - | - | - | - |

**KEY:** M1- M11 = eluted fractions of *M. inermis*
+ = present
- = absent

## 4.0 Conclusion

From the various parameters studied on the medicinal plants, *Lawsonia inermis* and *Mitragyna inermis,* the plants were found to be very rich in phytochemicals such as flavonoids, sterols, polyphenol, alkaloids and saponins in both plants, while anthraquinone was present in only *Lawsonia inermis.* The richness could justify the multiple therapeutic indications for which the plants are used for traditionally, for example, *Mitragyna inermis* is widely known for the treatment of fever, malaria, mental disorder, high blood pressure, dysentery, syphilis, wound, epilepsy and diabetes while *Lawsonia inermis*, popularly known as Henna is widely known for the treatment of fever, skin diseases, urinary tract infections, gonorrhea, jaundice, leprosy and also applied in the cosmetic industry for hair dyeing and body beautification.

